# Deoxycholic Acid (DCA) a Secondary Bile Acid, Mitigates Ovariectomy-Induced Bone Loss and promotes Bone Healing: A Novel Candidate for Orthopaedic Applications

**DOI:** 10.64898/2026.11.12.763604

**Authors:** Sumedha Yadav, Swati Rajput, Priyanca Ahlawat, Chaman Saini, Megha Sharma, Hanuman Prasad Sharma, Bhavuk Garg, Vikrant Manhas, Sahil Batra, Nupur Sharma, Jaswinder Singh Maras, Pradyumna K. Mishra, Divya Singh, Rupesh K. Srivastava

**Author notes:** **Send correspondence to:** Dr. Rupesh K. Srivastava, Additional Professor, Translational Immunology, Osteoimmunology & Immunoporosis Lab (TIOIL) An ICMR Collaborating Centre of Excellence on Bone Health Department of Biotechnology, All India Institute of Medical Sciences (AIIMS), New Delhi-110029, India and.

## Abstract

Postmenopausal osteoporosis (PMO) is characterized by accelerated bone loss, gut microbiota (GM) dysbiosis and alterations in microbial metabolites. Deoxycholic acid (DCA) is a secondary bile acid (BA) approved by the FDA for submental fat reduction, remains underexplored for its effects on bone health. This study investigated the association of DCA with PMO and its osteoprotective and bone regenerative potential. Integrated GM and plasma metabolomics analyses in ovariectomized (OVX) mice and clinical samples from PMO women revealed alterations in bacterial taxa associated with secondary BA metabolism, accompanied by reduced circulating DCA levels. Subsequently, DCA (20 mg/kg) administration for 45 days significantly improved trabecular and cortical bone microarchitecture in OVX mice, accompanied by suppressed osteoclastogenesis, enhanced osteoblastogenesis and modulation of circulating osteoclastogenic (IL-6, IL-9, IL-17A, and RANKL) and osteoblastogenic (OPG, TGF-β, IL-10, and IFN-γ) factors. Mechanistically, DCA suppressed osteoclast differentiation in peripheral blood mononuclear cells (PBMCs) from PMO women, reducing the expression of genes associated with osteoclast differentiation (NFATC1, NF-κB, TRAF6, TREM2, and SYNB), fusion (CSF1R, DC-STAMP, and CD47), and resorption (CTSK, ATP6V0D1, ATP6V0D2, CAR2, and SPP1). Furthermore, DCA promoted osteogenic differentiation and matrix mineralization in bone marrow-derived mesenchymal stem cells (BMSCs) from PMO women. In a femoral osteotomy model, 10-day DCA treatment enhanced calcium deposition at the fracture site and improved bone healing, accompanied by reduced osteoclastogenesis. Collectively, our findings identify reduced circulating DCA as a metabolic feature associated with PMO and demonstrate its osteoprotective and bone-regenerative potential, highlighting its promise for therapeutic repurposing in osteoporosis management and bone repair.

**Highlights:**

- OVX mice and PMO women have altered bacterial taxa associated with secondary BA metabolism.
- Circulating DCA levels are reduced in OVX mice and PMO women.
- DCA administration improves trabecular and cortical bone microarchitecture in OVX mice.
- DCA suppresses osteoclastogenesis and promotes osteoblastogenesis.
- DCA modulates bone-remodelling-associated cytokines and immune responses.
- DCA promotes bone regeneration and enhances fracture healing.

**Graphical Abstract:** 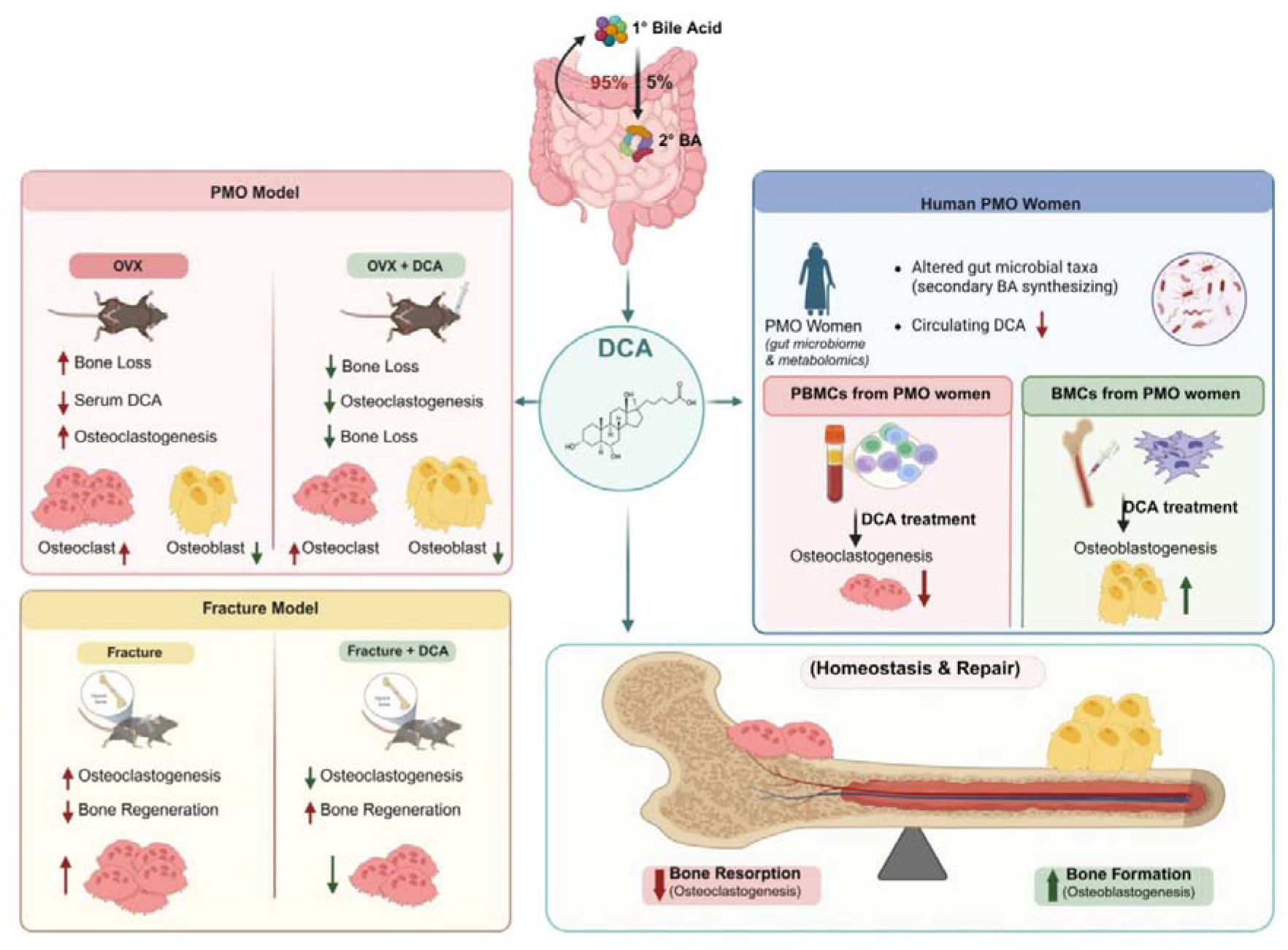

## Introduction

Osteoporosis is a systemic skeletal disorder, characterized by low bone mass and increased bone turnover leading to bone fragility and increased fracture risk [1]. It is the most common metabolic bone condition, affecting over 200 million people globally. It is widely acknowledged as a global public health concern that results in morbidity, mortality, and a strain on healthcare [2]. Postmenopausal osteoporosis (PMO) is the most prevalent type of osteoporosis in women, driven by estrogen deficiency following menopause. Low levels of estrogen are associated with increased osteoclast differentiation and activity while impairing osteoblast function, resulting in enhanced bone turnover with increased bone loss and an increased fracture risk. Estrogen regulates bone remodelling through the Wnt/β-catenin pathway and by regulating the RANKL/OPG ratio. Low estrogen levels lead to increased RANKL/OPG ratio thereby enhancing osteoclast differentiation and activity. Estrogen deficiency is also associated with increased pro-inflammatory cytokines such as TNF-α, IL-1β, and IL-6, which in turn are associated with enhanced bone loss [3–5]. Although, estrogen is a key determinant of PMO, increasing evidence indicates that bone remodelling is also regulated by other factors including, metabolic, immune, and endocrine signals originating beyond the skeletal microenvironment. Recent advances suggest the role of gut microbiota in modulating bone health both directly by regulating osteoclastogenesis and osteoblastogenesis, and indirectly by regulating immune system. These evidences led to growing interest in the “gut-bone” axis as an important contributor to skeletal homeostasis and the pathophysiology of osteoporosis [6,7].

The gastrointestinal tract is home to microbial ecosystem comprising over 100 trillion bacterial cells and over 3 million microbial genes. This microbial population is essential to the human body’s environmental homeostasis. Alterations in gut microbial composition have been reported in PMO women as well as in preclinical models of estrogen deficiency, impacting bone homeostasis through changes in intestinal permeability, inflammatory signalling, and microbial metabolic activity [8,9]. In particular, gut-associated metabolites (GAMs) provide a potential molecular link between microbial alterations and distant skeletal tissues. Microbial metabolites, including short-chain fatty acids (SCFAs), tryptophan-derived metabolites, and secondary bile acids (BAs), can modulate both immune and metabolic pathways involved in bone remodelling [10,11]. Among these, secondary BAs are of particular interest because their composition and biological activity are substantially shaped by the gut microbiome (GM). GM converts the primary BAs produced in the liver to the secondary BAs viz. lithocholic acid (LCA) and deoxycholic acid (DCA). DCA and LCA are signalling substances that further influence cellular differentiation, inflammation, and metabolic processes by activating the bile acid receptors G protein-coupled bile acid receptor 1 (TGR5) and farnesoid X receptor (FXR). These receptors are present in both bone-associated cells and can thus directly impact bone remodeling [12].

DCA has therefore emerged as a potentially relevant metabolite at the interface between microbial metabolism and skeletal regulation. Moreover, DCA is an FDA-approved drug for the treatment of submental fat in adults; however its effect on bone health remains underexplored [13]. Our previous study demonstrated that DCA attenuated lipopolysaccharide (LPS)-induced inflammatory bone loss, by attenuating osteoclastogenesis and enhancing osteoblastogenesis [14]. In the same model, DCA treatment was associated with restoration of LPS-induced GM alterations and increase of intestinal barrier integrity, including increased expression of the tight-junction proteins occludin and zonula occludens-1. Pharmacological inhibition of TGR5 or FXR further eliminated the protective effects of DCA, supporting the involvement of BA receptor signalling in its bone remodelling activity [14]. These findings thereby, suggest a functional connection between DCA, gut homeostasis, and bone remodelling under inflammatory conditions. Given that PMO represents a distinct form of bone loss driven predominantly by estrogen deficiency; these observations provided a rationale to investigate the relevance of DCA to bone health under estrogen deficient conditions.

Recent evidence has further implicated DCA in estrogen deficiency-associated bone loss; however, important aspects of the “gut-DCA-bone” relationship in human PMO remain unresolved. In particular, the relationship between alterations in the GM and circulating DCA in PMO women has not been comprehensively defined, and the functional effects of DCA on osteoclastogenesis and osteoblastogenesis in PMO women remain insufficiently characterized. Moreover, the potential role of DCA in bone regeneration, beyond its effect on bone remodelling remains to be determined.

In the present study, we investigated the role of DCA in estrogen deficiency-associated bone loss using complementary animal models and human samples. We first examined the GM composition and circulating DCA levels in both preclinical mice model and PMO subjects. Next, we evaluated the effects of exogenous DCA in ovariectomy-induced bone loss, with particular emphasis on osteoclastogenic and osteoblastogenic responses. Further, to establish the relevance of these findings to PMO, the effects of DCA was further examined in osteoclasts and bone marrow-derived mesenchymal stem cells (BMSCs) obtained from PMO subjects. Finally, using a femoral osteotomy model, we assessed whether DCA-associated effects on bone remodelling were also evident during skeletal repair. Together, these studies examine the “gut-DCA-bone” relationship and determine its effects on ovariectomy-induced bone loss, bone regeneration, and it’s potential in modulating bone remodeling cycle in PMO subjects.

## 2.0 Materials & Methods

### 2.1 Reagents and antibodies

Acid Phosphatase Leukocyte (TRAP) kit was purchased from Sigma (USA), while deoxycholic acid (DCA, CAS 83-44-3) was purchased from Cayman Chemical (USA). ProSpec (USA) provided the recombinant murine M-CSF (CYT-308) and RANKL (CYT-334). Gibco (USA) supplied culture media, such as RPMI-1640 and α-MEM, whereas tissue culture plasticware were purchased from Falcon (India). RT-PCR master mix was purchased from Promega (USA). The cDNA synthesis kit (1708890-Bio-Rad), gene-specific primers (*Ctsk*, *c-Fos*, Tgr5, *Fxr, Nfatc1*, tartrate-resistant acid phosphatase (trap), *Rankl*, *Runx2, Opg*, osteocalcin (*Ocn*), osterix (*Osx*), *CSF1R*, *DC-STAMP*, *CD47*, *NFATC1*, *NFκ-B*, *TRAF6*, *TREM*, *SYNB*, *ATP6VOD1*, *ATP6VOD2*, *CAR2*, and *SPP1* synthesised from Eurofins, India. Primer sequences were provided in **Table 1**, Alizarin Red S (A5533, Sigma, USA). ELISA kits were purchased from Elabscience (China). Plastic wares were purchased from Tarsons (India).

### 2.2 Postmenopausal Osteoporotic (OVX) mice model

8–10-week-old female C57BL/6 mice were used for ex vivo and in vivo studies. The animals were housed in the animal facility of the All India Institute of Medical Sciences (AIIMS), New Delhi, India, under specified pathogen-free (SPF) conditions and provided with ad libitum access to autoclaved food and drinking water. For the *in vivo* experiments, mice were randomly assigned to one of three groups: sham, ovariectomized (OVX), and OVX+DCA. In the sham mice, the ovaries were intact; in OVX and OVX+DCA mice, the ovaries were removed bilaterally after anaesthetizing with ketamine (70-80 mg/kg) and xylazine (10 mg/kg). Mice in the OVX+DCA group were orally administered with 20 mg/kg of DCA for 45 days starting a week after surgery. At the end of the experimental period (45 days), mice were euthanized, and blood, bones and bone marrow, were collected for downstream analysis. All procedures were carried out with due approval from the Institutional Animal Ethics Committee of AIIMS (338/IAEC-1/202), New Delhi, India. A part of the approved protocol was also published previously [14].

### 2.3 Micro-CT

After dissection, the hind limbs from all groups were collected, cleared of surrounding muscles, and fixed in 4% paraformaldehyde. Micro-CT scanning and analysis were performed in SkyScan 1076 scanner (Aartselaar, Belgium). The samples were properly positioned in the sample holder and scanned at 50 kV and 204 mA using a 0.5-mm aluminium filter. The reconstructed images were generated using NRecon software. For trabecular bone analysis, 100 slices of the region of interest (ROI) were drawn from the secondary spongiosa. CTAn software was used for 3D-analysis of the trabecular region to determine bone parameters including, bone volume/tissue volume (BV/TV), trabecular number (Tb. N), trabecular thickness (Tb. Th), trabecular separation (Tb. Sp). Cortical parameters were assessed using 2D analysis and included tissue area (T. Ar), tissue perimeter (T. Pm), bone perimeter (B. Pm), and crossectional thickness (Cs. Th). The BMD of the femur and tibia was calculated from the volume of interest defined for the trabecular and cortical regions. Hydroxyapatite phantom rods (4 mm diameter) with known BMDs of 0.25 and 0.75 g/cm³ were used for calibration.

### 2.4 Osteoclast Differentiation and TRAP Staining

Bone marrow cells (BMCs) were isolated from the femurs and tibias of mice by flushing the bones with α-MEM containing 10% heat-inactivated FBS. Following red blood cell lysis using 1× RBC lysis buffer, cells were cultured overnight in T-25 flasks containing endotoxin-free α-MEM supplemented with 10% heat-inactivated FBS and M-CSF (35 ng/mL). After 16-24 hours, non-adherent cells were collected, and 5 × 10^4^ cells were seeded into 96-well plates containing α-MEM supplemented with M-CSF (30 ng/mL) and RANKL (50 ng/mL). On day 3, half of the culture medium was replaced with fresh α-MEM supplemented with the respective factors.

On day 4, osteoclast differentiation was assessed using TRAP staining according to the manufacturer’s protocol. Briefly, cells were washed with PBS and fixed for 10 min at room temperature in a solution of citrate, acetone, and 3.7% formaldehyde, followed by two washes with PBS. Cells were then incubated with TRAP substrate solution at 37°C in the dark for 5-15 min. Multinucleated TRAP-positive cells containing three or more nuclei were considered osteoclasts. Images were acquired using inverted microscopes (Eclipse TS100, Nikon; EVOS, Thermo Fisher Scientific, USA) and osteoclast surface area was quantified with ImageJ software (NIH, USA).

### 2.5 Quantitative PCR (q-PCR)

To isolate RNA from mice bone marrow cells and human PBMC-derived osteoclasts, the RNeasy Mini Kit (Qiagen, USA) was used in accordance with the manufacturer’s instructions. Bio Analyser (Agilent Technologies, Singapore) was used to assess the integrity of RNA (28 S and 18 S), and only samples with RNA Integration Number (RIN) ≥7.0 were included in further analysis. Nanodrop spectrophotometer (Nanodrop Technologies, USA) was used to determine the concentration of RNA, and 260/280 from 1.8 to 2.0 was considered to be optimum. For cDNA synthesis Thermo Scientific cDNA synthesis kit was used. Customised primers c-FOS, TRAP, CTSK, NFATC1, RANKL, OPG, OSTERIX, OSTEOCALCIN, RUNX2, FXR, TGR5, CSF1R, DC-STAMP, CD47, NFKΒ, TRAF6, TREM2, SYNB, ATP6VOD1, ATP6VOD2, CAR2, and SPP1 were used for gene specific analysis. For each reaction, 25 ng of cDNA was used in each well along with primers and 2X SYBR green PCR master mixes (Promega, USA). Threshold cycle data of each gene were normalised with GAPDH and expressed in relative gene expression using the 2^−ΔΔCT^ method.

### 2.6 Enzyme-Linked Immunosorbent Assay

Serum levels of IL-6, IL-9, IL-17A, RANKL, OPG, TGF-β, IL-10, and IFN-γ, were measured using commercially available ELISA kits, following the previously published protocols [15].

### 2.7 Osteoblastogenesis and ALP staining

BMCs were isolated from the femur and tibia of 8-12-week-old C57BL/6J mice by flushing with α-MEM containing 10% heat-inactivated FBS. The cells were washed three times with α-MEM and seeded in 96-well plates. The media was replenished with 50% fresh media on day 1 and 80% on day 2, respectively. On day 3, the complete media was replenished with media containing osteogenic factors, ascorbic acid (50 μg/ml), and β-glycerophosphate (10 mM). Thereafter, fifty percent of the media was replenished every third day with fresh osteogenic media.

On day 14, ALP staining was performed. For ALP staining, cells were washed with PBS and fixed in 4% paraformaldehyde for 30 min at room temperature. Following fixation, the cells were washed and incubated with BCIP/NBT substrate solution at 37°C in the dark for 15 min [16]. The reaction was stopped by washing with PBS. Stained cultures were imaged using an EVOS XL Core microscope (Thermo Fisher Scientific, USA). ALP staining intensity was quantified using ImageJ software (NIH, USA).

### 2.8 Study Cohort

The study was designed as a cross-sectional case-control study. We recruited post-menopausal women, and classified as either control with T-score > −1 and osteoporosis with T-score ≤ −2.5 determined by non-invasive Dual Energy X-ray Absorptiometry (DEXA) (**Table 2**). PMO women on any anti-osteoporotic medications, undergoing any form of treatment (comorbidities) or on antimicrobial therapy were excluded from the study. This protocol was approved by Institute Ethics Committee, All India Institute of Medical Sciences with reference number AIIMSA2259/06.09.2024, RP-47/2024. Informed written consent was taken from all participants before sample collection. The bone marrow samples were collected from the participants admitted for knee replacement surgery and sample were collected during surgery.

### 2.9 Untargeted Metabolomics

5 mL of blood samples were withdrawn from the recruited PMO subjects and plasma was separated using blood and Lymphoprep™ (StemCell, Norway) (3:1) density gradient centrifugation at 800 g for 25 min at 0/0 acceleration/deceleration. Isolated plasma samples were subjected to untargeted metabolomics of control (n=20) and PMO (*n* = 57) using the below mentioned protocol.

An organic phase extraction protocol was used to isolate metabolites from 200 µl plasma sample. About 400 µl methanol was added to plasma samples and kept overnight at −20°C for protein precipitation. These samples were centrifuged at 13,000 rpm for 10 min. The protein pellet was discarded, and the supernatant was isolated, dried under vacuum conditions and reconstituted with (90:5:5) 90% water:5% internal standards:5% acetonitrile and then loaded for reverse-phase chromatography in a C18 column (Thermo Scientific™25003102130: 3 μm, 2.1 mm, and 100 mm) using an ultra-high-performance liquid chromatographic system followed by high-resolution orbitrap mass spectrometry (HRMS). All the samples were pooled together and spiked with internal standards (Dinose b: 1mg/mL; MCPA: 1mg/mL; Dimetrazole: 1mg/mL) and external standards (Cholesterol: 0.1mg/mL; Colchicine: 4mg/mL; Imparinine: 2mg/mL; Roxithromycin: 2mg/mL; Amiloride: 1mg/mL; Atropine:2mg/mL) and analysed in serial dilutions of 1:1,1:2,1:4 and 1:8 for determining analytical reproducibility. Further, Compound Discoverer 3.0 was used to identify metabolite features (ThermoFisher Scientific, Waltham, United States). The features were annotated using mass list, Chemspider, mz Vault, Metabolika and mzCloud™. Identified and annotated features were subjected to log normalization and Pareto scaling using a MetaboAnalyst 6.0 (http://metaboanalyst.ca.) server and used to perform principal component analysis and partial least square discriminant analysis, volcano plots, and others.

### 2.10 Targeted Metabolomics

The quantification of deoxycholic acid in patient plasma samples and animal sera samples utilized liquid chromatography coupled with mass spectrometry (LC-MS/MS). The analysis was conducted using the 6500+ triple quadrupole mass spectrometer with a linear ion trap (Sciex, MA, USA). The mass spectrometer was integrated with Exion ultra high-performance liquid chromatography (UHPLC). The UHPLC equipped itself with an online degasser, a binary pump, a computer-controlled sample cooler, and a column heater. All parameters of the UHPLC and mass spectrometer were configured using Analyst (version 1.7.1; Sciex, MA, USA). The metabolites were chromatographically separated on a HypersilGold C18 column (50×2.1 mm, 1.9 µm; ThermoScientific, USA). A gradient approach was employed in LC-based analysis, utilizing ultra-pure water with 0.1% formic acid (A) and acetonitrile:methanol (2:1) with 0.1% formic acid (B). The baseline condition was initiated with 50% mobile phase A and maintained for the next 0.5 min. The concentration of A was reduced to 28% in the next 6 min. The mobile phase A was further reduced to 20% in next 8 minutes and further to zero after 10 seconds. The condition was held for one minute and forty seconds and then returned to the baseline level after 1.5 minutes. The overall duration of the analysis was 18 minutes, and the injection volume was 5 µL. The column and sample tray oven were maintained at 40°C and 10°C, respectively. Probenecid was used as an internal standard (IS) for the analysis. A multiple reaction monitoring (MRM) mode was used for the quantitative analysis of the samples using a Turbo Ion Spray (ESI) source operated in negative ionization mode. The MRM transition for DCA and probenecid were m/z 391.3->345.3/327.3 and 284.1->240.1, respectively. All the samples were thawed at room temperature. Samples were vortexed carefully and centrifuged briefly. A 12.5 µL sample was taken and extracted with 150 µL of extraction solvent (acetonitrile containing 100 ng/mL probenecid). The mixture was incubated at 4 °C for 30 min at 600 rpm on a temperature-controlled mixer (ThermoMixer C, Eppendorf, USA). Next tubes were centrifuged at 7820 g for 10 minutes at 4°C. The clear supernatant was transferred to a new tube and evaporated to dryness using a CentriVap Cold Trap vacuum concentrator (Labconco, Kansas City, Missouri, USA). The dried samples were finally reconstituted in methanol:water (1:1) and analyzed by LC-MS/MS (Sciex, MA, USA).

### 2.11 Osteoclastogenesis in human PBMCs

10 ml of blood was withdrawn and layered with Ficoll in a 1:3 (1 Ficoll, 3 blood) ratio to generate a density gradient. Centrifuged the layered blood at 800g for 20 minutes at room temperature, with breaks set at 0. After centrifugation, the white layer of mononuclear cells containing lymphocytes was collected. The cells were washed thrice with 1X PBS at 4°C. The first wash was at 400 g for 12 minutes, followed by 300 g for 10 minutes and a 3^rd^ wash at 200 g for 8 minutes. The pellet collected was resuspended in α-MEM media, and cells were counted. 1X10^6^ cells/well of a 96-well plate were seeded in the α-MEM media. Cells were incubated for 2 hours at 37°C to let the cells adhere and then washed thrice with plain media to remove suspension or loosely adhered cells. After washing, adherent cells were treated with α-MEM supplemented with MCSF (25 ng/ml) and RANKL (100 ng/ml) in the presence or absence of DCA at 5µM concentration. Fifty percent of the media was replenished every 3^rd^ day, and TRAP staining was performed on day 14 as described in section 2.4 [17].

### 2.12 Osteoblastogenesis in Human BMSCs and ALP activity

Bone marrow samples were obtained from the PMO participants undergoing knee replacement. The whole bone marrow sample was cultured in Dulbecco’s Modified Eagle Medium (DMEM) low glucose media. After 48 hours, the media was replenished to remove non-adherent cells, and the cells were allowed to grow till 80% confluency was attained with intermittent media change. At 80% confluency, cells were passaged and grown till passage 3. After passage 3, the plastic-adherent cells were obtained, and flow cytometry analysis was done to get the mesenchymal stem cells of more than 95% purity. The MSCs were further used for osteoblastogenesis till passage 7.

5000 cells per well of a 96-well plate were seeded and incubated for 24 hours in DMEM low glucose media. After 24 hours, the media was removed and replaced with fresh media containing osteogenic factors (ascorbic acid, 50 μg/ml, and β-glycerophosphate, 10 mM). MSCs were treated with or without DCA at a concentration of 5 μM. The cells were cultured for 14 days and on day 14 ALP activity were performed.

For the ALP activity assay, the media was aspirated, and the cells were washed with PBS. The cells were freeze-thawed thrice for cell lysis. To achieve this, the plate was dried and kept at −20°C for 2 hours, and once frozen, it was kept at 37°C for 30 minutes for thawing. The freeze-thaw process was repeated thrice. Next, p-nitrophenyl phosphate (pNPP) was added to the wells, and once the yellow color was developed, the plate was read at 405 nm using a NanoDrop spectrophotometer/fluorimeter (BioTek Synergy, USA) [18,19].

### 2.13 Alizarin Red S staining

BMSCs were cultured as mentioned in Section 2.12 in the presence or absence of DCA at a concentration of 5 μM in complete media supplemented with osteogenic factors. Media was replenished every 3rd day for the 21 days. On day 21, alizarin red S staining was performed. For ARS staining, culture medium was aspirated, and cells were washed twice with PBS. Cells were then fixed with 10% formaldehyde (100 µL per well) for 30 min at room temperature, followed by two additional PBS washes. A 2% alizarin red solution prepared in distilled water (50 µL per well) was added, and plates were incubated at 37°C until calcium deposits were visibly stained. To quantify mineralization, the bound dye was eluted with 100 µL of a solution containing 10% acetic acid, 20% methanol, and 70% distilled water, followed by incubation for 15 min. Absorbance was recorded at 450 nm using a BioTek Synergy H1 (Agilent Technologies, USA) spectrophotometer/fluorimeter [20]

### 2.14 Osteotomy model

C57BL/6 male mice aged 8-10 weeks were used to study the bone regeneration potential of DCA in a drill hole-induced fracture model in an *in vivo* study. The animals were kept in the animal facility of the All India Institute of Medical Sciences (AIIMS), New Delhi, India, under specified pathogen-free (SPF) conditions. For the *in vivo* experiments, mice were randomly assigned to one of two groups: Fracture and Fracture + DCA. Briefly, a standardized cortical defect was created in the mid-diaphysis of the right femur by using a 25-gauge needle (∼0.5 mm diameter) inserted from the anterior to the posterior aspect of the femur. Following surgery, mice were administered orally with DCA (20 mg/kg) for 10 days to assess its effect on early bone repair. At the end of the experiment (day 11), mice were sacrificed, and the fractured bones were collected. All procedures were carried out with due approval from the Institutional Animal Ethics Committee of AIIMS (338/IAEC-1/2022), New Delhi, India.

### 2.15 Calcein labelling

In the femur osteotomy model, mice received a subcutaneous injection of calcein (20 mg/kg, dissolved in PBS) 24 hours before the sacrifice. The bones were then harvested and subjected to micro-CT analysis. The bones were embedded in the acrylic resin and subsequently, sectioned into 60 µm using an Isomet slow-speed precision cutter. The sections were visualised under a fluorescence microscope (Eclipse Ti, Nikon), and images were captured. Fluorescence intensity was quantified using ImageJ software (NIH, USA).

### 2.16 F-Actin ring formation assay

Osteoclast cytoskeletal organization was evaluated using phalloidin staining. BMCs were isolated from both Fracture and Fracture + DCA treated mice and cultured in 96-well plates, and on day 4, cells were processed for F-actin visualisation. Cultures were rinsed twice with PBS, fixed in 4% paraformaldehyde for 20 min, and permeabilised with 0.1% Triton X-100 for 5 min. To reduce nonspecific staining, cells were incubated with 1% BSA for 30 min before labelling with FITC-conjugated phalloidin for 30 min at room temperature in the dark. Nuclear counterstaining was performed with DAPI (1 µg/mL, 5 min, dark). F-actin ring structures were examined under a fluorescence microscope (Eclipse Ti, Nikon) [21].

### 2.17 16S rRNA sequencing data analysis

16S rRNA sequencing analysis was performed using our previous data deposited in the Mendeley data repository (https://data.mendeley.com/research-data/?query=10.17632/7j57nk95sv.1). The raw data were further processed for quality filtering and trimming. Next the data was deionised, followed by chimera removal using DADA2 pipeline. SILVA reference database was used to identify high-quality amplicon sequence variants (ASVs). The ASVs abundance table, taxonomic assignments, and sample metadata were generated using the phyloseq package in R programming. The differential abundance of various genus & species were plotted in form of heatmap.

### 2.18 Statistical analysis

All experiments were independently repeated using different biological replicates to ensure the reproducibility of the results. The data are displayed as mean ± SEM. One-way ANOVA followed by Dunnett’s multiple-comparisons test was used to compare more than two groups, while Student’s t-test (paired or unpaired, as appropriate) was used for comparisons between two groups. p < 0.05 (*), p < 0.01 (**), p < 0.001 (***), and p < 0.0001 (****). The statistics are explained in depth in the figure legends.

## Results

### 3.1 Ovariectomy leads to gut dysbiosis in mice

To investigate the association between estrogen deficiency and composition of secondary BA producing bacteria in the gut, we first performed 16S rRNA sequencing of both Sham and ovariectomized (OVX) mice. **Fig. 1A and Fig. S1**, shows the analysis of microbial composition at the genus level and we observed that the genus *Bacteroidota* were reduced and *Firmicutes* to *Bacteriodata* ratio were significantly enhanced in ovariectomized mice compared with the sham mice. Moreover, species level analysis revealed differences in the relative abundance of several bacterial taxa between the two groups (**Fig. 1B**). Notably, OVX mice exhibited significant reduction in the abundance of various species including *Bacteroides acidifaciens* (p < 0.001), *Bacteroides caecimuris* (p < 0.0001), *Lactobacillus gasseri* (p < 0.05) and *Ligilactobacillus* (p < 0.001) compared to sham mice (**Fig. 1C**). Further to validate our data we next performed secondary analysis of an independently generated mouse 16S rRNA sequencing dataset comprising Sham and ovariectomized (OVX) mice [22]. Similar to our results, this dataset too produced results showing phylum and genus level differences in the relative abundance of several bacterial taxa between the two groups **(Fig. S1)**. Notably, OVX mice exhibited significant reduction in the abundance of various genera. The relative abundance of *Clostridia_UCG-014* (p < 0.01) was significantly reduced in OVX mice compared to sham. However, the relative abundance of Bifidobacterium was non-significant. Moreover, the relative abundance of *Ligilactobacillus* (p < 0.05), *Lactobacillus* (p < 0.01) and *Bacteroides* (p < 0.05) was significantly reduced in OVX mice compared to sham **(Fig. S1)**. This clearly suggests that ovariectomy is associated with alterations in the gut microbial composition. Importantly, the overall microbial alterations identified in our experimental cohort were concordant with that observed in the independent secondary dataset. The reduced relative abundance of *Ligilactobacillus, Lactobacillus,* and *Bacteroides* related taxa across the two datasets supports the association of ovariectomy with specific alterations in GM composition. Moreover, these microbiota possess bile salt hydrolase enzyme activity, which is majorly associated with secondary BA synthesis. [23–25].

**Fig. 1:**
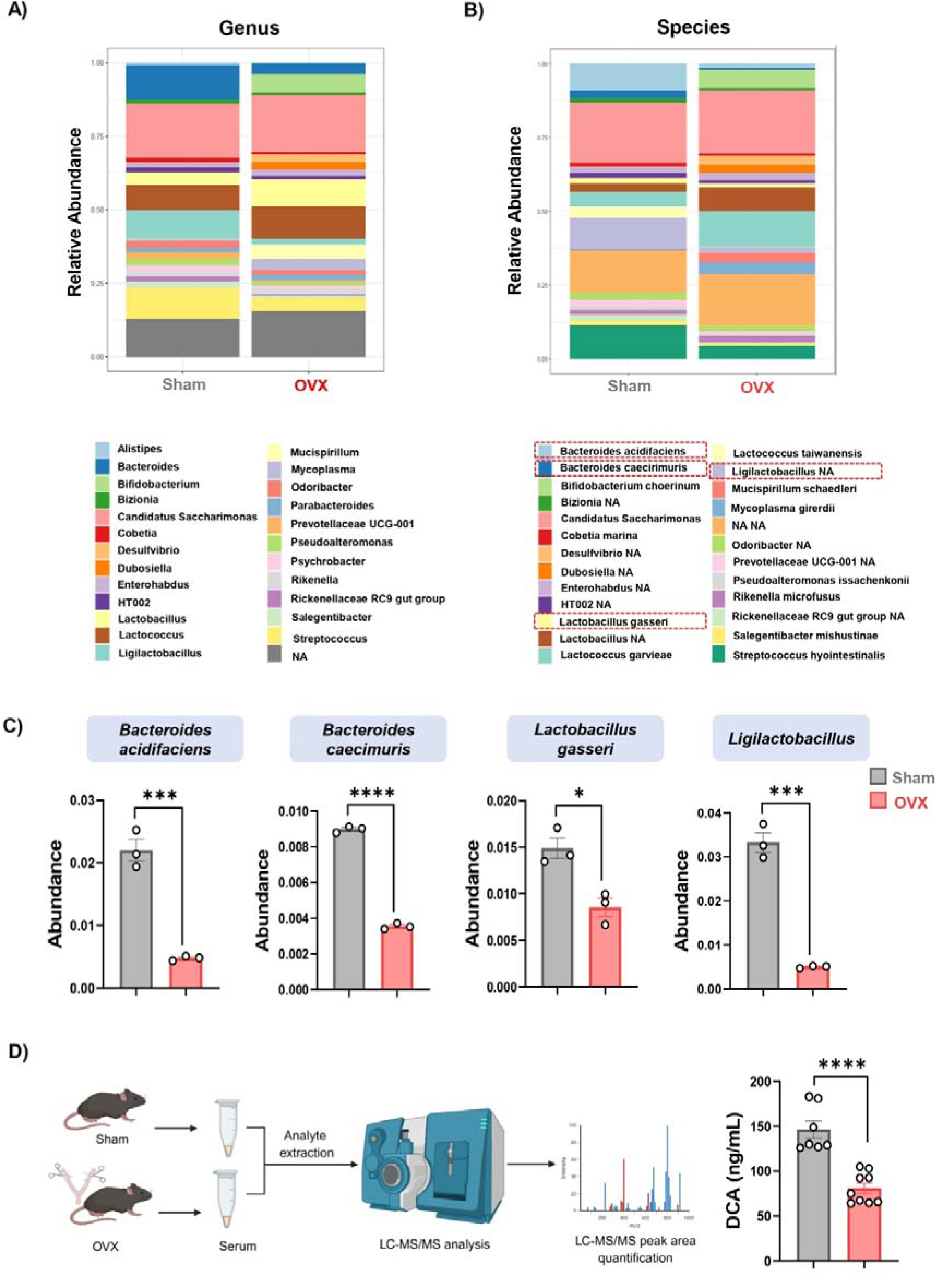
Ovariectomy leads to gut dysbiosis in mice. Heatmap showing differential gut microbiota composition among Sham and OVX groups at **(A)** genus level, **B)** species level. **C)** Plots representing the GM composition at species level (n=3/group). **D)** DCA quantification in mice sera in different groups (n=7-10/gp). Data are represented as mean ± SEM. Statistical analyses were performed using unpaired student t-test. * p < 0.05, ** p < 0.01, *** p < 0.001, and **** p < 0.0001. DCA: Deoxycholic acid

Further, to check whether these alterations in GM are translated in postbiotics too, specifically the secondary BAs, we next performed mass spectrometry analysis of serum obtained from sham and OVX mice. Interestingly, we found that the concentration of DCA in OVX mice was significantly (p< 0.0001) reduced in the serum as compared to the sham mice (**Fig. 1D**). These findings suggests that decreased secondary BA synthesizing bacteria in OVX-mice play a crucial role in modulating bone health. This led us further to investigate how DCA modulates bone health.

### 3.2 DCA attenuated OVX-induced bone loss *in vivo*

To investigate the osteoprotective effects of DCA under PMO conditions, we first developed an OVX-induced PMO model in C57BL/6J mice (**Fig. 2A**). The µCT analysis revealed impaired trabecular bone microarchitecture of the femur and tibia in OVX mice compared with sham. The reduced bone mineral density (BMD), lower bone volume fraction (BV/TV), decreased trabecular number (Tb. N), decreased trabecular thickness (Tb. Th) and a pronounced increase in trabecular separation (Tb. Sp) were observed in OVX mice. Notably, DCA treatment attenuated these pathological alterations, restored bone microarchitecture and improved histo-morphometric parameters (**Fig. 2B-E**). Additionally, DCA treatment also significantly attenuated cortical bone loss in both femur and tibia, marked by tissue area (T. Ar), tissue perimeter (T. Pm), cross-sectional thickness (Cs. Th), endosteal perimeter (E. Pm) and medullary area (M. Ar) in both femur and tibial cortical bones (**Fig. 3A-D**). Altogether, our *in vivo* results indicate that DCA effectively ameliorates bone loss under pre-clinical postmenopausal conditions. DCA exhibits a strong ability to prevent OVX-induced skeletal deterioration by maintaining both trabecular and cortical bone microarchitecture, highlighting its therapeutic potential for postmenopausal osteoporosis.

**Fig. 2:**
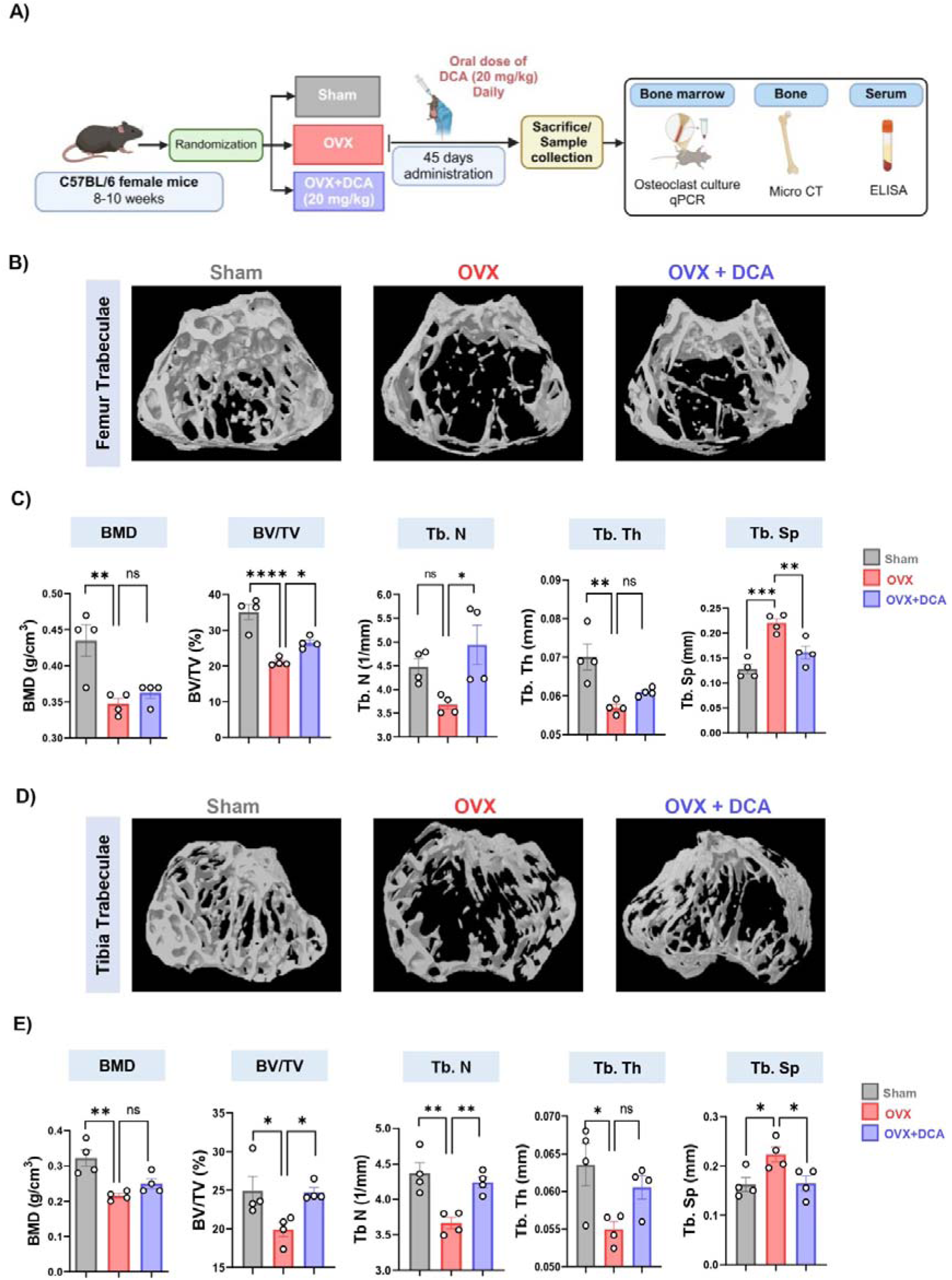
DCA administration improves trabecular bone microarchitecture in OVX mice. **A)** Schematic representation of the *in vivo* experimental design showing the groups included: Sham, OVX, and OVX + DCA. **B)** 3D µCT reconstruction of femur trabecular bone. **C)** Plots representing the histomorphometric parameters of femur trabecular bone, including BMD, BV/TV, Tb. N, Tb. Th, and Tb. Sp. **D)** 3D representative images of tibia trabecular bone. **E)** Bar graphs showing the histomorphometric parameters of tibia trabecular bone including BMD, BV/TV, Tb. N, Tb. Th, and Tb. Sp. Data are represented as mean ± SEM (n = 4). Statistical analyses were performed using one-way ANOVA followed by Dunnett’s test. * p < 0.05, ** p < 0.01, *** p < 0.001 and **** p < 0.0001. DCA: Deoxycholic acid; BMD: Bone mineral density; BV/TV: Bone volume/ tissue volume; Tb. N: Trabecular number; Tb. Th: Trabecular thickness; Tb. Sp: Trabecular separation

**Fig. 3:**
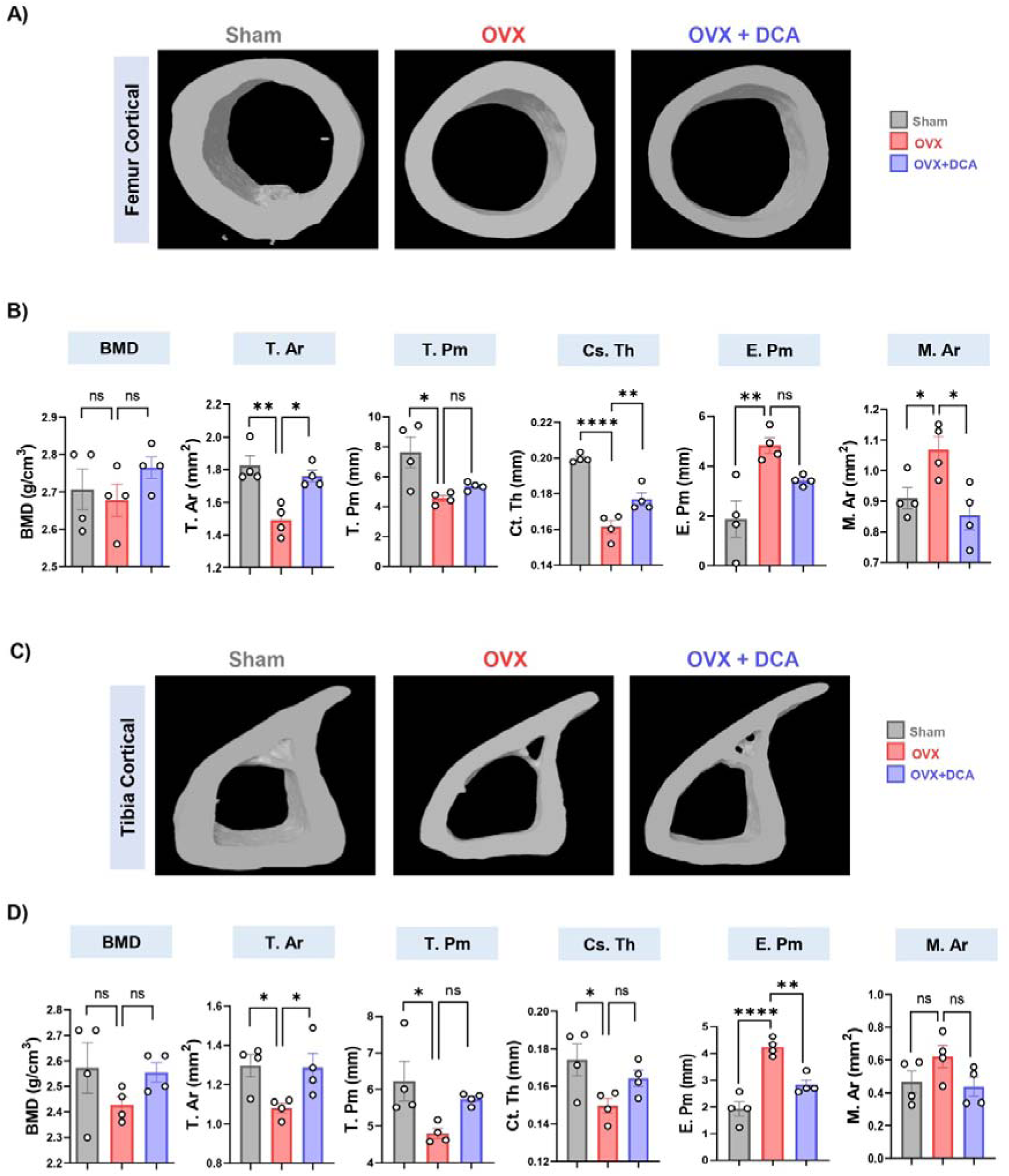
DCA administration improves cortical bone parameters in OVX mice. **A)** 3D µCT reconstruction of femur cortical. **B)** Plots representing the histomorphometric parameters of femur cortical bone, including BMD, T. Ar, T. Pm, Ct. Th, E. Pm and M. Ar **D)** 3D representative images of tibia cortical bone. **E)** Bar graphs showing the histomorphometric parameters of tibia cortical bone including BMD, T. Ar, T. Pm, Ct. Th, E. Pm and M. Ar. Data are represented as mean ± SEM (n = 4). Statistical analyses were performed using one-way ANOVA followed by Dunnett’s test. * p < 0.05, ** p < 0.01, *** p < 0.001 and **** p < 0.0001. DCA: Deoxycholic acid; BMD: Bone mineral density; T. Ar: Tissue area; T. Pm: Tissue perimeter; Ct. Th: Crossectional thickness; E. Pm: Endosteal perimeter; M. Ar: Medullary area

### 3.3 DCA suppressed Osteoclastogenesis in OVX mice

Further, to investigate how DCA treatment affects osteoclast differentiation in OVX mice, we performed osteoclastogenesis of bone marrow cells obtained from sham, OVX and OVX + DCA mice. TRAP staining analysis showed that the TRAP positive cells, osteoclast cells with >3 nuclei and area of osteoclast were significantly enhanced (p < 0.001) in OVX compared to the sham mice. Interestingly, DCA treatment significantly reduced the TRAP positive cells and cells with >3 nuclei as well as the area of osteoclast (p < 0.0001) (**Fig. 4A-D)**.

**Fig. 4:**
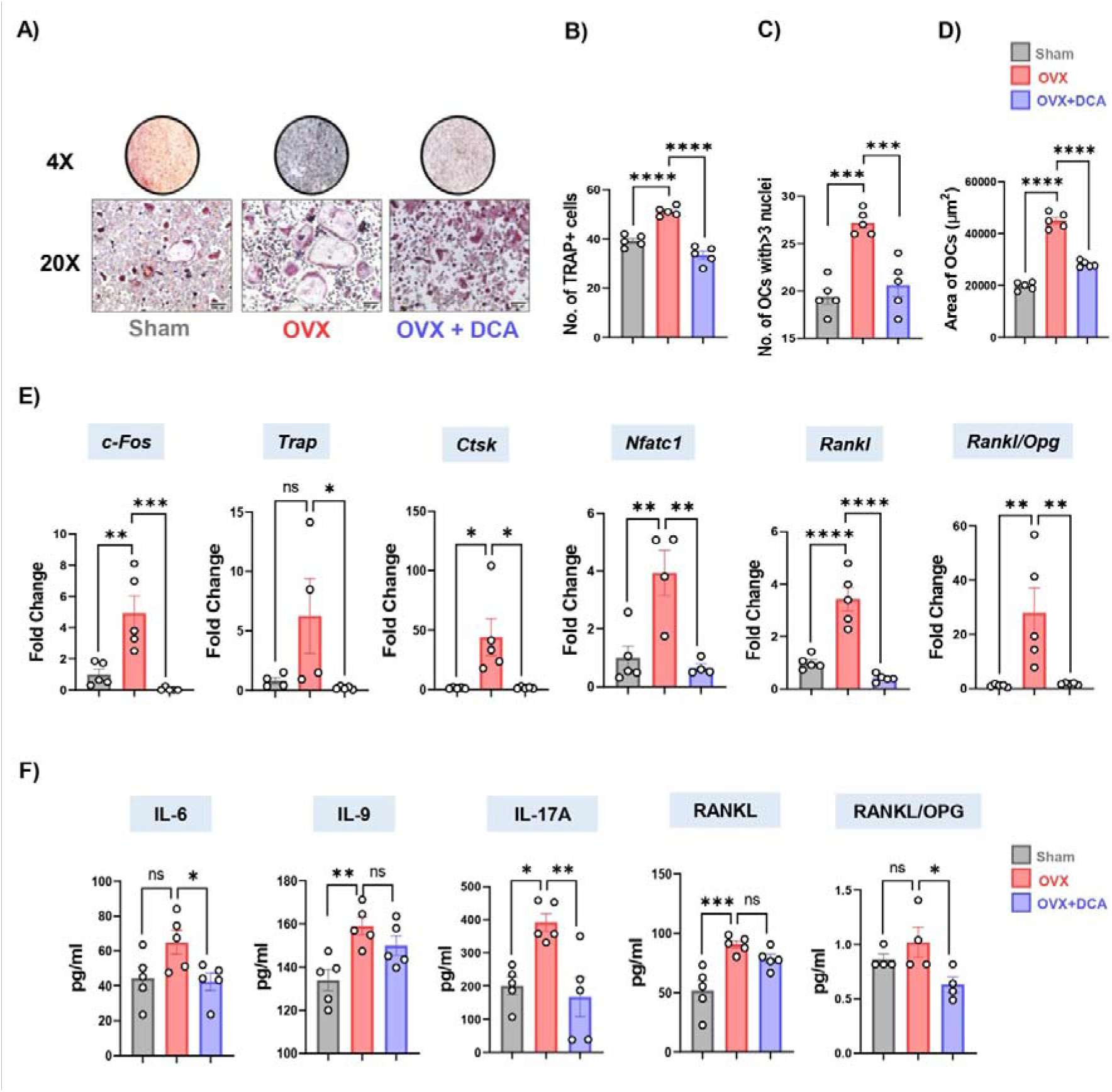
DCA administration prevents osteoclastogenesis. **A)** TRAP-stained osteoclasts cultured from the bone marrow of Sham, OVX and OVX + DCA mice at 20X (scale bar, 500 µm). **B)** Bar graph representing TRAP-positive cells. **C)** Number of multinucleated osteoclast cells. **D)** Area of osteoclasts. **E)** mRNA expression of osteoclastogenic genes, including *c-Fos*, *Trap, Ctsk, Nfatc1, Trap, Rankl and Rankl/Opg ratio*. GAPDH was used as a normalising control. **F)** Serum levels of osteoclastogenic factors (IL-6, IL-9, IL-17A, RANKL, and RANKL/OPG) in Sham, OVX and OVX + DCA. Data are expressed as mean ± SEM (n=4-5). Statistical analyses were performed using one-way ANOVA followed by Dunnett’s test. * p < 0.05, ** p < 0.01, *** p < 0.001 and **** p < 0.0001. DCA: Deoxycholic acid

To further understand the molecular basis of these cellular alterations, we studied the expression of key osteoclastogenic genes in BMCs of sham, OVX and OVX + DCA. Consistent with the above results, OVX mice showed that the osteoclastogenic markers, *c- Fos*, Trap, *Ctsk, Nfatc1,* and *Rankl* were significantly enhanced (**Fig. 4E**). The DCA treatment markedly restored the alteration in osteoclastogenic markers caused by ovariectomy (**Fig. 4E**). Moreover, the osteoclastogenic factors viz. IL-6 (p < 0.05), IL-9, IL-17A (p < 0.01), and RANKL (p < 0.05) were significantly reduced by DCA treatment which was significantly high in OVX mice (**Fig. 4F**). Moreover, the higher RANKL/OPG ratio in OVX mice at the RNA & protein level was significantly reduced by DCA treatment (p < 0.01) (**Fig. 4E-F**). These results suggest that DCA inhibits osteoclastogenesis and thereby, prevents bone loss in OVX mice.

### 3.4 DCA promotes osteoblastogenesis in OVX mice

Next, to ascertain the osteoblastogenic potential of DCA treatment in OVX mice, we performed osteoblastogenesis in BMCs collected from sham, OVX and OVX + DCA mice. **Fig. 5A** represents the experimental layout of osteoblast culture. ALP staining revealed that DCA treatment significantly enhanced osteoblastogenesis (p < 0.05) which was reduced in OVX mice, further ALP quantification, confirmed these results (**Fig. 5B-C**). Concordant with the ALP results, DCA treatment non-significantly enhanced the expression levels of key osteoblastogenic markers including, *Opg, Osterix, Osteocalcin* and *Runx2* in OVX mice (**Fig. 5D**). DCA further enhanced the expression of *Fxr* (p < 0.01) and *Tgr5*; the bile acid receptors which were reduced in OVX mice (**Fig. 5D**). Further, DCA restored the key osteoblastogenic factors including, OPG, TGF-β (p < 0.001), IL-10 (p < 0.01), and IFN-γ (p < 0.01) which were significantly reduced in OVX mice (**Fig. 5E**). Altogether, our results suggest that DCA prevents bone loss in PMO model by its dual modulatory effect on both osteoclastogenesis as well as osteoblastogenesis.

**Fig. 5:**
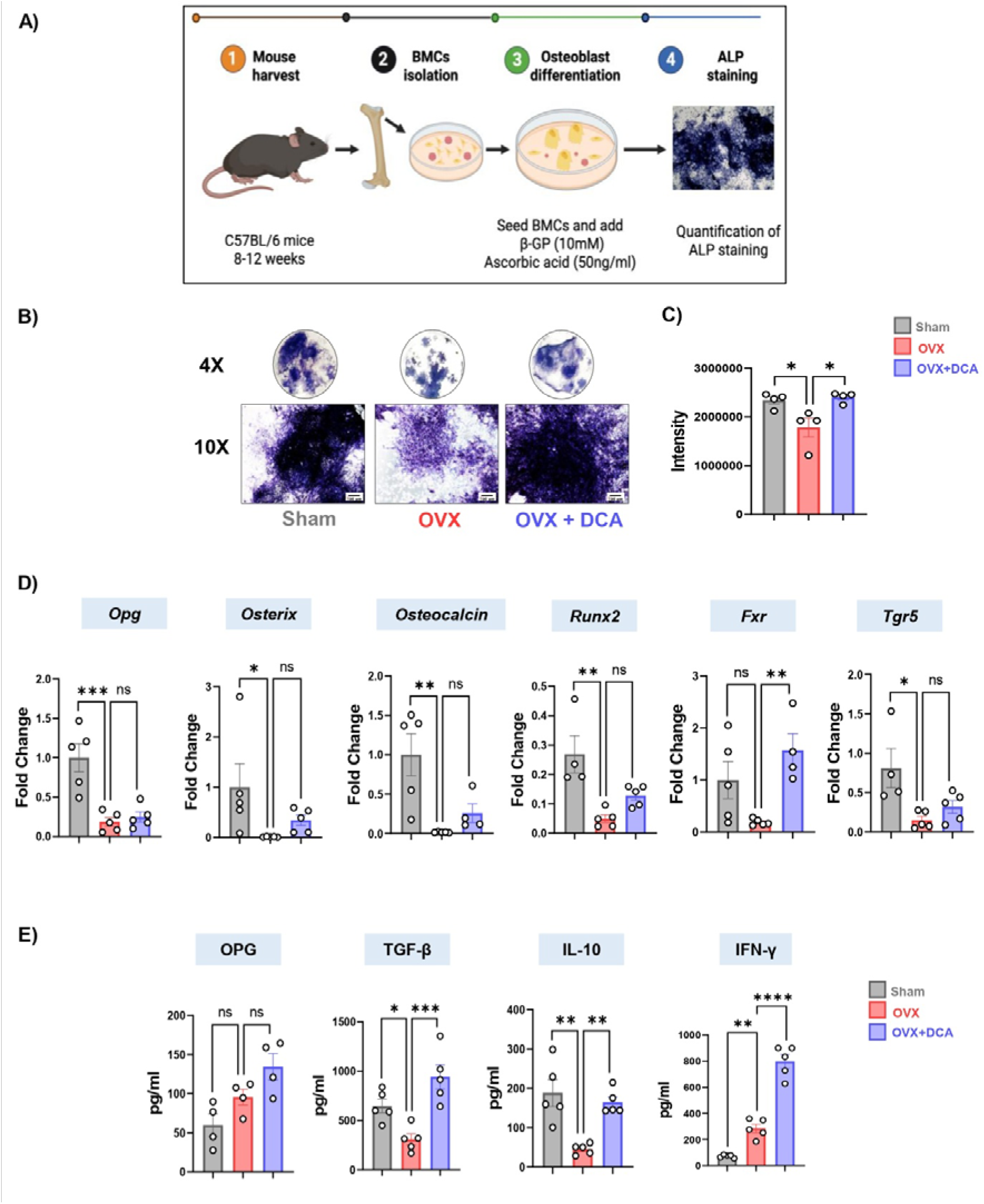
DCA enhances osteoblastogenesis. **A)** Schematic representation of osteoblast culture from Sham, OVX and OVX + DCA at 10X (scale bar 100 µm). **B)** ALP-stained osteoblasts cultured from the bone marrow of Sham, OVX and DCA-treated OVX mice. **C)** Bar graph representing intensity of ALP stain. **D)** mRNA expression of anti-osteoclastogenic genes, *Opg, osterix, osteocalcin, Runx2*, and BA receptors *Fxr* and *Tgr5*. GAPDH was used as a normalising control. **E)** Serum levels of anti-osteoclastogenic factors (OPG, TGF-β, IL-10, and IFN-γ) in Sham, OVX and OVX + DCA. Data are expressed as mean ± SEM (n = 4-5). Statistical analyses were performed using one-way ANOVA followed by Dunnett’s test. * p < 0.05, ** p < 0.01, *** p < 0.001 and **** p < 0.0001. DCA: Deoxycholic acid

### 3.5 DCA producing GM and circulating DCA levels were decreased in PMO subjects

Our previous results from the pre-clinical study demonstrated that the secondary BA synthesising bacteria were decreased in OVX mice along with reduced concentration of DCA in serum. Further, DCA treatment attenuated the bone loss occurred in OVX mice, along with inhibiting osteoclastogenic genes and promoting osteoblastogenic gene expression. Next, we were interested to find out whether similar pattern is observed in the human samples. To investigate the alteration in human GM, secondary analysis of an independent dataset was performed [26]. Interestingly, we found that the secondary BAs synthesising bacteria *Bifidobacterium adolescentis* (p < 0.05)*, Bacteroides vulgatus* (p < 0.0001)*, Bacteroides massiliensis* (p < 0.001)*, Bacteroides caccae* (p < 0.0001) and *Parabacteroides goldstenii* (p < 0.01) were significantly reduced (**Fig. 6A**), confirming the pattern similar to that observed in ovx mice.

**Fig. 6:**
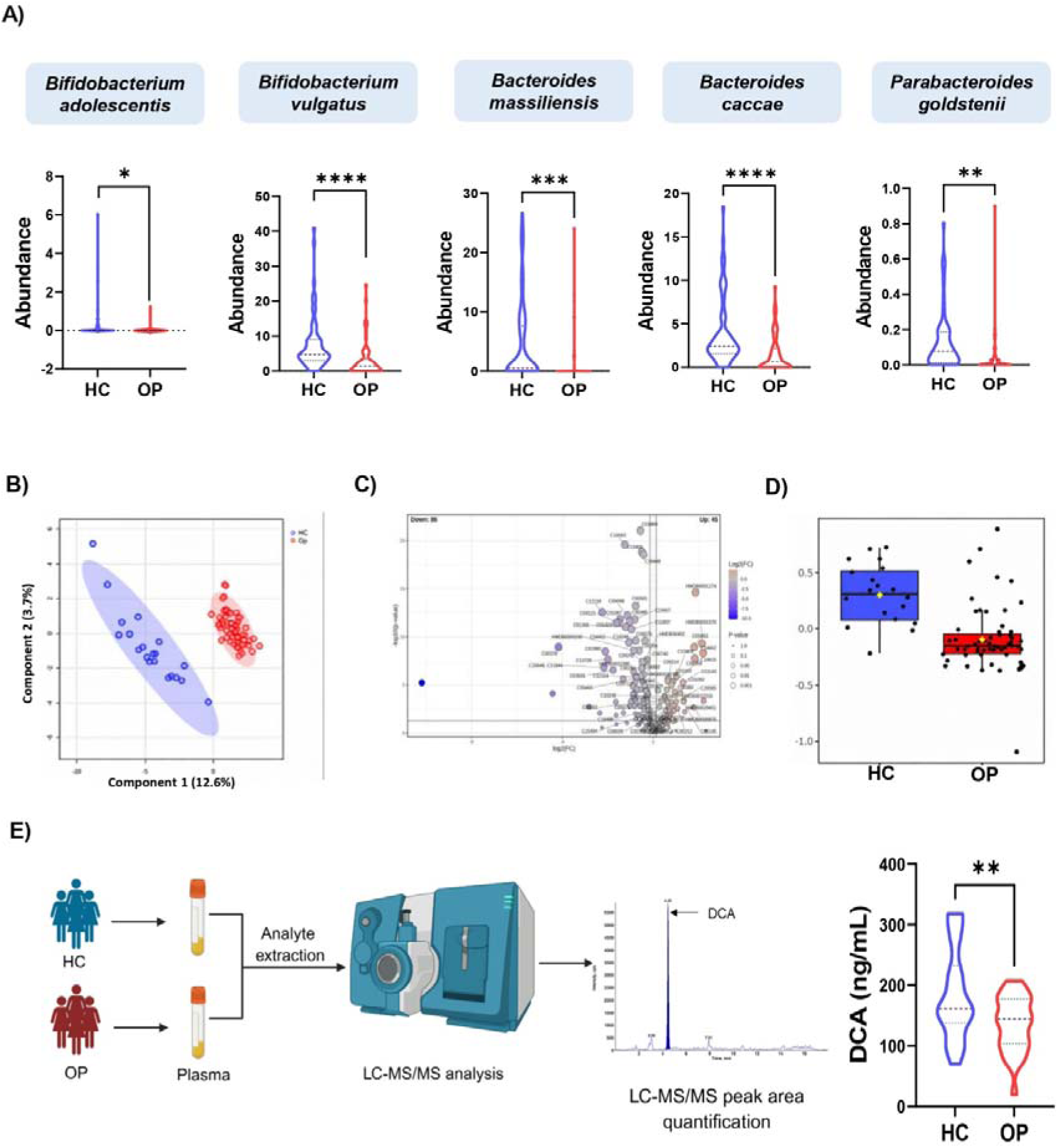
Postmenopausal Osteoporosis leads to Dysbiosis. **A)** Plots showing GM composition in healthy control (HC) and postmenopausal osteoporotic women (OP) at species level (n = 60). **B-D)** PLS-DA plot, volcano plot & Box plot showing downregulated DCA (C04483) in plasma untargeted metabolomics profile of HC =20 and OP=57. **E)** Targeted metabolomics of human plasma for DCA quantification in HC (n=20) and OP (n=30). Data are represented as mean ± SEM. Statistical analyses were performed using unpaired student t-test. * p < 0.05, ** p < 0.01, *** p < 0.001 and **** p < 0.0001. DCA: Deoxycholic acid

Further to validate the role of secondary BAs in modulating bone health, we performed untargeted metabolomic profile of 84 postmenopausal women (control = 20, osteoporotic = 57) and our results showed down regulation of DCA (Kegg ID-C04483) in osteoporotic samples (FC-1.1; p≤0.05) (**Fig. 6B-D**). Taking lead from these findings, LC-MS/MS based targeted quantification of DCA in plasma samples of the same cohort clearly demonstrated significantly lower levels in postmenopausal osteoporotic women (p < 0.01) (**Fig. 6E**). These findings clearly indicated a persistent role of DCA in the establishment and progression of postmenopausal osteoporosis. Thus, therapeutic potential of DCA as a bone remodeling modulator in estrogen-deficient conditions is strongly supported by the above findings.

### 3.6 DCA suppressed Osteoclastogenesis and enhanced Osteoblastogenesis in PMO subjects

Next, we wanted to validate whether our findings could be translated in human samples, and for the same we cultured PBMCs obtained from PMO women in osteoclastogenic media with or without DCA. **Fig. 7A** depicts the experimental design. TRAP staining revealed that osteoclastogenesis was significantly inhibited by DCA treatment in PBMCs of osteoporotic women (p < 0.01) (**Fig. 7B**). Number of osteoclast with more than 3 nuclei and area of osteoclast were significantly reduced (P < 0.01) by DCA (**Fig. 7C-D**). Further, to validate these findings at molecular level we performed real time PCR to investigate the expression levels of key osteoclastogenic genes. We observed that DCA treatment significantly suppressed the expression of osteoclast fusion-associated genes, including, *CSF1R* (p < 0.05), *DC-STAMP* (p < 0.05) and *CD47* (p < 0.05) compared to control (M-CSF + RANKL; MR) (**Fig. 7E**). DCA also suppressed key osteoclast differentiation genes *Nfatc1* (p < 0.001)*, NF-κB* (p < 0.05)*, TRAF6* (p < 0.001) and signalling genes *TREM2* (p < 0.01) and *SYNB* (**Fig. 7F-G**). Notably, DCA significantly reduced the expression of bone-resorption associated genes, including, *CTSK (*p < 0.05)*, ATP6VOD1 (*p < 0.05)*, ATP6VOD2 (*p < 0.05)*, CAR2 (*p < 0.05) and *SPP1* p < 0.01 (**Fig. 7H**).

**Fig. 7:**
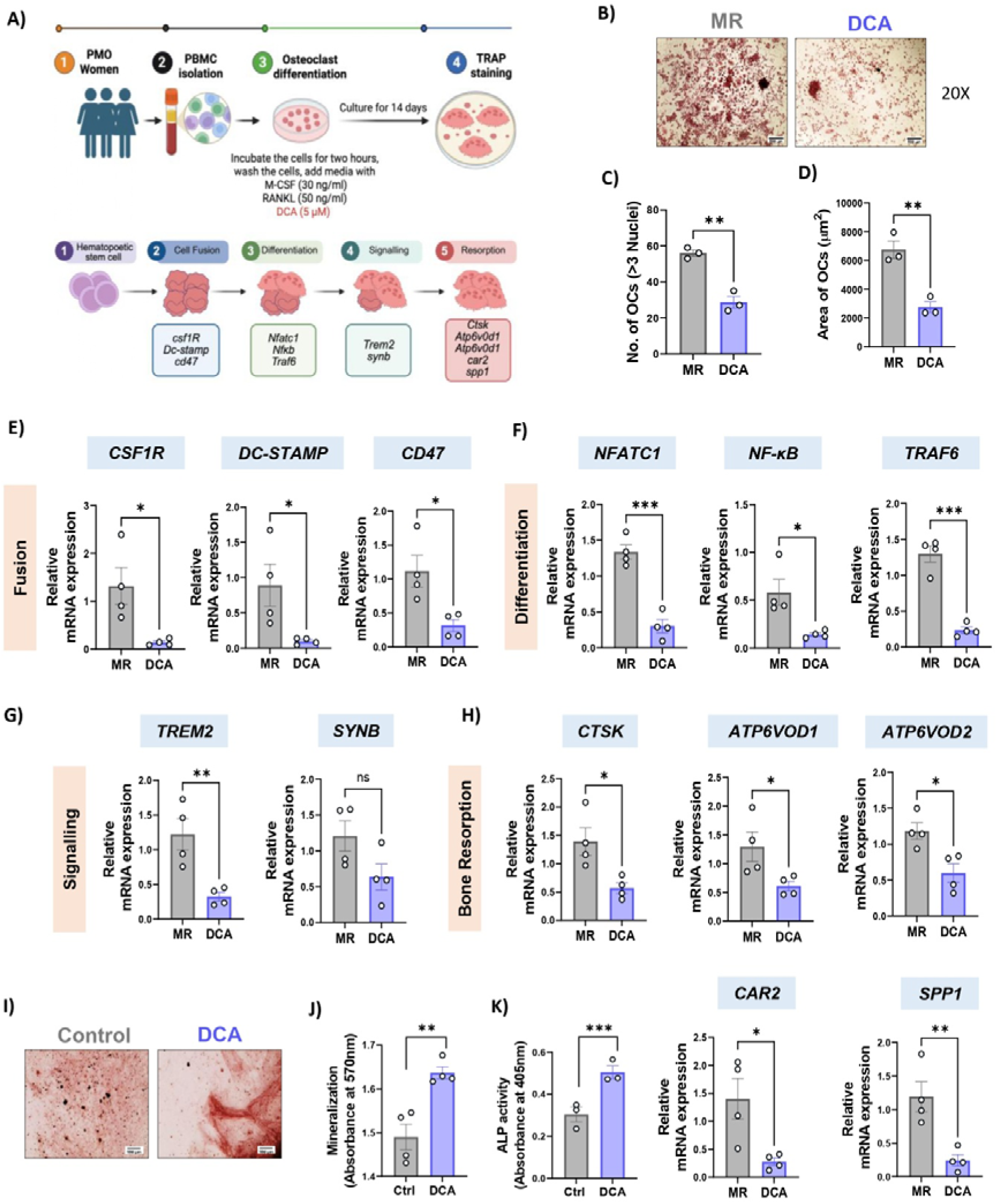
DCA suppressed Osteoclastogenesis and enhanced Osteoblastogenesis in PMO subjects. **A)** Schematic representation of osteoclast culture from human PBMCs. **B)** Representative images of TRAP-stained osteoclasts in control & DCA-treated PBMCs at 20X (scale bar, 500 µm). **C)** Quantitative analysis of number of multinucleated osteoclast cells. **D)** Area of osteoclasts. Bar graph representing qPCR analysis of osteoclastogenic genes: **E)** Fusion, **F)** Differentiation, **G)** Signalling, and **H)** Bone resorption in osteoclast culture of human PBMCs in presence or absence of DCA. **I)** Alizarin red staining (ARS) on mineralised nodules formed from MSCs cultured with or without DCA for 21 days, images were taken at 4X (scale bar, 100 µm). **J)** Bar graphs showing the quantification of ARS. **K)** Bar graph showing ALP activity of MSCs cultured with or without DCA for 14 days. Data were presented as mean ± SEM (n=3-4). Statistical analyses were performed using the paired student t-test. * p < 0.05, ** p < 0.01, *** p < 0.001 and **** p < 0.0001. Ctrl: Control, DCA: Deoxycholic acid, MR: M-CSF and RANKL

Further, to investigate the osteoblastogenic potential of DCA in PMO subjects, firstly, we checked the purity of human MSCs via flow cytometry and our results have shown that these MSCs were found to be positive for CD90, CD105, CD73, > 90% and negative for CD45 (**Fig. S2**). Next these human MSCs were treated with DCA for their effect on ALP & ARS staining. We found a significant increase in ALP activity in the DCA treated osteoblasts (p < 0.001). In consistence with the ALP activity ARS staining showed significant increase in the formation of calcium nodules in MSCs obtained from PMO subjects (p < 0.01). Further, quantitative analysis of ARS staining confirmed our above findings and revealed that DCA treatment significantly enhanced matrix mineralization as compared to the control group in human MSCs (**Fig. 7I-K**). These results indicate that DCA promotes early as well as late osteoblast differentiation. Collectively, these findings suggest that DCA significantly inhibits osteoclastogenesis and enhances osteoblastogenesis in PMO subjects. DCA exerts its anti- osteoclastogenic effect by inhibiting multiple molecular mechanisms of osteoclastogenesis by encompassing osteoclast fusion, differentiation, signalling and resorption in osteoclast culture of human PBMCs. Moreover, DCA further enhances the osteoblastogenic potential of MSCs by promoting early osteoblastogenic differentiation and late stage matrix mineralization.

### 3.7 DCA enhanced bone healing at fracture site

Following the beneficial effects of DCA on bone homeostasis in preclinical PMO model and its regulation of osteoclast and osteoblast activity in PMO subjects, next we investigated whether DCA could also promote bone regeneration in fracture. As osteoporosis is the primary cause of fracture, thereby, investigating, how DCA is impacting bone healing following fracture becomes important. To investigate the effect of DCA on fracture healing, we developed a femoral osteotomy model in C57BL/6J mice and treated them with DCA (**Fig. 8A** represents the experimental layout). Micro-CT analysis of the fractured bones obtained from control (untreated) and DCA treated mice showed a marked improvement in bone regeneration at the osteotomy site in DCA treated mice (**Fig. 8B**). Additionally, BMD and BV/TV at the injury site were also significantly (p < 0.05) enhanced following DCA treatment, indicating enhanced mineralization during the healing process (**Fig. 8C**).

**Fig. 8:**
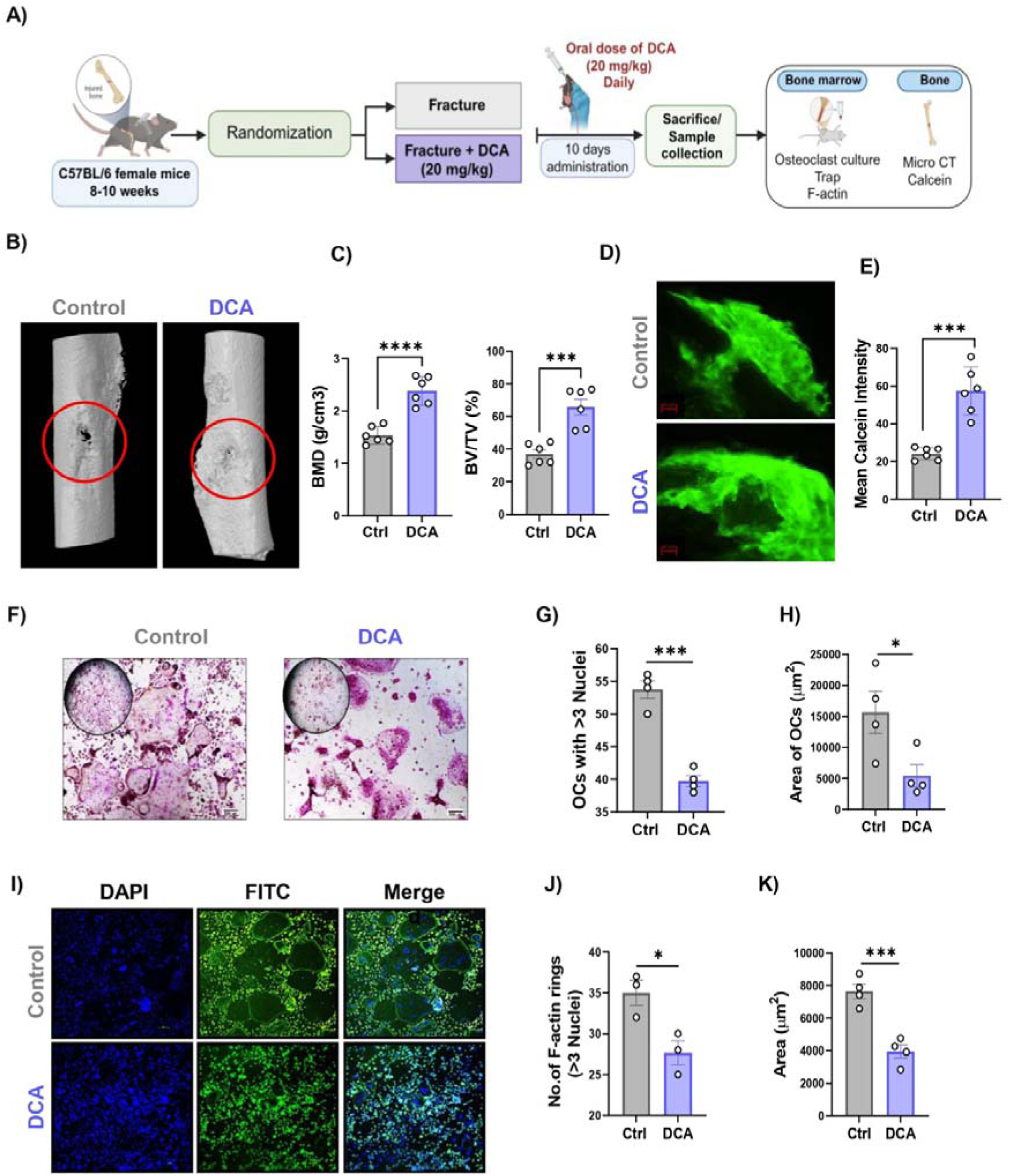
DCA administration accelerates fracture healing. **A)** Schematic representation of the *in vivo* experimental design showing the groups included: Fracture and Fracture + DCA (n=6/group). **B)** Representative 3D μCT images of callus at the osteotomy site. **C)** Quantification of BMD and BV/TV. **D)** Representative image of calcein at osteotomy site. **E)** Quantification of calcein intensity in control and DCA treated group. **F)** TRAP-stained osteoclasts cultured from the bone marrow of control and DCA-treated mice. **G)** Bar graph number of multinucleated osteoclast cells. **H)** Area of osteoclasts. **I)** F-actin and nuclei stained with phalloidin and DAPI, respectively. **J)** Bar graph representing the number of multinucleated cells. **K)** Area of F-actin rings. Data are expressed as mean ± SEM (n = 3). Statistical analyses were performed using the paired student t-test.* p < 0.05, ** p < 0.01, *** p < 0.001 and **** p < 0.0001. Ctrl: Control, DCA: Deoxycholic acid

Further, to assess bone regeneration at the fracture site, new bone formation at the fracture site was evaluated using calcein fluorochrome labelling. Calcein labelling enables visualization of actively mineralizing surfaces and thus serves as an indicator of newly formed bone. Fluorescence imaging and mean intensity of the fracture callus revealed enhanced calcein incorporation in the DCA-treated group compared with the control group (**Fig. 8D-E**), indicating increased mineral deposition and new bone formation at the site of injury. These observations were consistent with the micro-CT findings, thereby further supporting the role of DCA in promoting bone regeneration during fracture healing.

To further evaluate the effect of DCA on osteoclast differentiation during fracture healing, bone marrow cells isolated from the Fracture (control) and Fracture + DCA groups were cultured in osteoclastogenic medium supplemented with M-CSF and RANKL. TRAP staining performed on day 4 demonstrated a marked reduction in TRAP-positive multinucleated osteoclasts with more than 3 nuclei in the DCA-treated group compared with the control group (p < 0.001) (**Fig. 8F-G**). In addition, DCA treatment significantly decreased the overall area of multinucleated osteoclasts; indicating reduced osteoclast maturation and resorptive potential (p < 0.05) (**Fig. 8H**).

Furthermore, to evaluate the effect of DCA on osteoclast functional activity in fracture mice, F-actin ring formation was examined (a cytoskeletal structure essential for osteoclast- mediated bone resorption). F-actin staining revealed that DCA treatment markedly reduced both the number (p < 0.05) and size of F-actin rings compared with the control group (p < 0.001) (**Fig. 8I-K**). The presence of well-organized F-actin rings is a hallmark of actively resorbing osteoclasts; therefore, these findings clearly indicate that DCA compromises the functional activity of osteoclasts in addition to inhibiting their differentiation. Overall, our findings indicate that DCA has the potential to promote bone regeneration by suppressing both the formation and resorptive function of osteoclasts during fracture healing.

## 4. Discussion

Postmenopausal osteoporosis (PMO), a manifestation of ageing and estrogen deficiency, disrupts the balance between bone formation and bone resorption, ultimately leading to enhanced bone loss. Although the effects of estrogen deficiency in the skeleton deterioration are well established, growing evidence suggests that alterations in the GM and its metabolites further contribute to bone homeostasis, influencing osteoporosis progression. In the present study, we identified DCA, a secondary bile acid, as a potential metabolic mediator linking estrogen deficiency-associated gut dysbiosis with altered bone remodelling. Our findings revealed that OVX was accompanied by changes in GM composition and a marked reduction in circulating DCA levels. Importantly, similar alterations in GM composition and reduced DCA levels were further observed in PMO women, highlighting the clinical relevance of these findings. Moreover, DCA supplementation attenuated OVX-induced deterioration of bone microarchitecture and modulated bone remodelling by suppressing osteoclastogenesis, while promoting osteoblastogenesis. Consistent with these *in vivo* observations, DCA suppressed osteoclastogenesis and promoted osteoblastogenesis in human cells derived from PMO subjects. These findings further support its potential role in regulating bone homeostasis.

The alterations in GM composition observed in OVX mice are consistent with growing evidence that the “gut-bone” axis is affected by estrogen deficiency-associated bone loss. Several bacterial taxa, including *Bacteroides*, *Lactobacillus*, and *Ligilactobacillus*, were reduced in OVX mice, with comparable changes observed in an independent mouse dataset. Consistent with these findings, GM analysis of PMO women revealed a significant reduction in several bacteria associated with secondary BA metabolism, including *Bifidobacterium adolescentis*, *Bacteroides vulgatus*, *Bacteroides massiliensis*, *Bacteroides caccae*, and *Parabacteroides goldstenii*. Among these, *B. adolescentis* and *B. vulgatus* have been reported to possess 7α-dehydroxylation activity, contributing to the conversion of primary BAs into secondary BAs [24]. Similarly, *B. caccae* has been associated with secondary BA-producing potential [25], whereas *P. goldstenii* possesses hydroxysteroid dehydrogenase activity and contributes to secondary BA metabolism [23]. Although, secondary BA biosynthesis involves multiple microbial and host-mediated steps and the observed taxonomic alterations do not establish direct DCA production by individual taxa, these alterations may reflect changes in the overall microbial capacity for BA metabolism. In this context, the concomitant reduction in circulating DCA levels in both OVX mice and PMO women suggests a possible association between estrogen deficiency-associated GM alterations and a diminished systemic secondary BA pool. These findings are consistent with earlier reports [27] demonstrating that OVX-induced dysbiosis is associated with alteration in bone remodelling. Furthermore, our previous study demonstrated that DCA attenuated LPS-induced inflammatory bone loss and restored GM balance, along with suppressing osteoclastogenesis and promoting osteoblastogenesis through FXR and TGR5 receptors [14].

To investigate the therapeutic potential of DCA under estrogen-deficient conditions, we further evaluated its effects on bone microarchitecture and bone remodelling in OVX mice. Micro-CT analysis revealed that DCA treatment significantly improved bone microarchitecture in both trabecular and cortical bone compartments. These structural improvements were accompanied by suppression of osteoclast differentiation in BMCs, along with reduced expression of key osteoclastogenic genes, including *c-Fos*, *Rankl, Ctsk, Nfatc1*, and *Trap,* and a reduced *Rankl/Opg* ratio. Additionally, DCA treatment suppressed inflammatory and osteoclastogenic cytokines, while restoring anti-inflammatory and osteoblastogenic cytokine levels in the serum of OVX mice. Consistent with these findings, DCA treatment markedly attenuated osteoclastogenesis in PBMCs derived from PMO women. This effect was associated with reduced expression of genes involved in osteoclast precursor survival, fusion, and differentiation, including *CSF1R*, *DC-STAMP, CD47, NFATC1, NF-κB,* and *TRAF6*. Furthermore, the downregulation of *CTSK, ATP6V0D1, ATP6V0D2, CAR2,* and *SPP1* indicates reduced expression of genes associated with osteoclast function and resorption. Together, these findings extend our previous observations on the osteoprotective effects of DCA and highlight its potential to modulate osteoclast- associated pathways under estrogen-deficient conditions. Thus, the findings from OVX mice and PMO-derived PBMCs provide complementary evidence supporting the anti- osteoclastogenic effects of DCA.

These findings further support the emerging role of BA receptor signalling in the regulation of bone remodelling. The increased expression of BA receptors FXR and TGR5 following DCA treatment in OVX mice suggests that these signalling pathways may contribute to the observed skeletal benefits. Previous studies have demonstrated that TGR5 deficiency enhances osteoclastogenesis and exacerbates OVX-induced bone loss, whereas TGR5 activation suppresses osteoclast differentiation [28]. Similarly, FXR has been implicated in the regulation of both osteoblast and osteoclast activity, with its activation favouring osteoblast differentiation and limiting osteoclastogenesis [29]. In agreement with these findings, our previous study demonstrated that pharmacological inhibition of TGR5 or FXR attenuated the anti-osteoclastogenic effects of DCA in inflammatory bone loss [14]. Collectively, these observations suggest that the osteoprotective effects of DCA under estrogen-deficient conditions may be mediated, at least in part, through the activation of FXR- and TGR5-associated signalling pathways.

In addition to its anti-osteoclastogenic effects, DCA also promoted osteogenic differentiation in human mesenchymal stem cells (MSCs), as demonstrated by enhanced ALP activity and ARS staining. These findings provide functional evidence of the osteoblastogenic potential of DCA in human cells. Interestingly, this functional response was observed despite the lack of significant restoration of osteoblastogenic gene expression, including *Opg, Osx, Ocn,* and *Runx2*, in the bone marrow of OVX mice. This suggests that the osteoblastogenic effects of DCA may not be fully reflected by changes in steady-state transcriptional markers in vivo. This observation is consistent with previous evidence implicating FXR and TGR5 in osteoblast regulation, wherein FXR activation promotes Runx2-associated osteoblast differentiation and TGR5 signalling enhances osteoblastic differentiation and mineralization [30]. Collectively, these findings suggest the pivotal role of DCA in regulating bone remodelling through coordinated effects on both osteoclast and osteoblast cells.

Beyond its effects on estrogen deficiency-associated bone loss, DCA further demonstrated bone regenerative potential in the femoral osteotomy model. Improved BMD and BV/TV at the osteotomy site, along with enhanced nascent calcium deposition, suggest increased mineral deposition during bone repair in DCA-treated mice. In parallel, DCA reduced TRAP- positive multinucleated osteoclasts and F-actin ring formation in BMCs derived from fractured bone marrow, demonstrating attenuation of both osteoclast differentiation and functional activity. The concomitant increase in mineral deposition and reduction in osteoclast activity suggest that DCA may modulate the cellular balance underlying bone remodelling during repair. Altogther, these findings extend the skeletal effects of DCA beyond the prevention of bone loss under estrogen-deficient conditions and further support its potential role in promoting bone regeneration.

The present study has certain limitations. Firstly, the potential adverse effects of DCA were not evaluated. Furthermore, the effects of DCA on bone mechanical strength and bone turnover indicators, including bone stiffness, maximum load, energy absorption, and biochemical markers of bone formation and resorption, remain to be investigated. Further mechanistic and translational studies are warranted to establish its efficacy and clinical applicability, particularly to understand its long-term effects on bone quality, remodelling, and fracture resistance. Despite these limitations, our findings highlight DCA as a novel therapeutic candidate for improving bone health under estrogen-deficient conditions. Importantly, it’s osteogenic and bone regenerative potential suggests that its therapeutic relevance may extend beyond the prevention of bone loss to promoting bone repair and regeneration. The fact that DCA is already an FDA-approved drug for reducing submental fat (double chin), the present findings provide a basis for exploring its repurposing potential in bone health. Further investigations will thus be essential to establish its safety, efficacy, and translational applicability in the management of osteoporosis.

## Acknowledgment

SY, SR, CS, PA, MS and RKS acknowledge the Department of Biotechnology, AIIMS, New Delhi, India, for providing infrastructural facilities and centralized Core Research Facility (CCRF) of AIIMS, New Delhi. SY, thank DBT for the research fellowship.

## Data availability

All data generated or analysed during this study are included in this published article and its supplementary information files. The current study’s data are available from the corresponding author upon reasonable request.

## Credit authorship contribution statement

RKS contributed to the conception and design of the study; SY contributed to the acquisition and analysis of data; SY contributed to drafting the text or preparing the figures. SR did ALP activity and ARS staining and helped with drafting. CS did qPCR, PA and HPS did the targeted serum metabolomics, PA, NS, and JSM did the untargeted plasma metabolomics and MS helped during *in vivo* studies. DS helped in MicroCT analysis, PKM provided valuable inputs. BG VM and SB provided patient samples. All authors reviewed the manuscript. All authors contributed to the article and approved the submitted version.

## Declaration of generative AI use

During the drafting of this publication, the authors used AI to help with language refinement, grammar checking, and all content was carefully reviewed and approved by the authors, who bear full responsibility for the final product.

## Supplementary

**Fig. S1:**
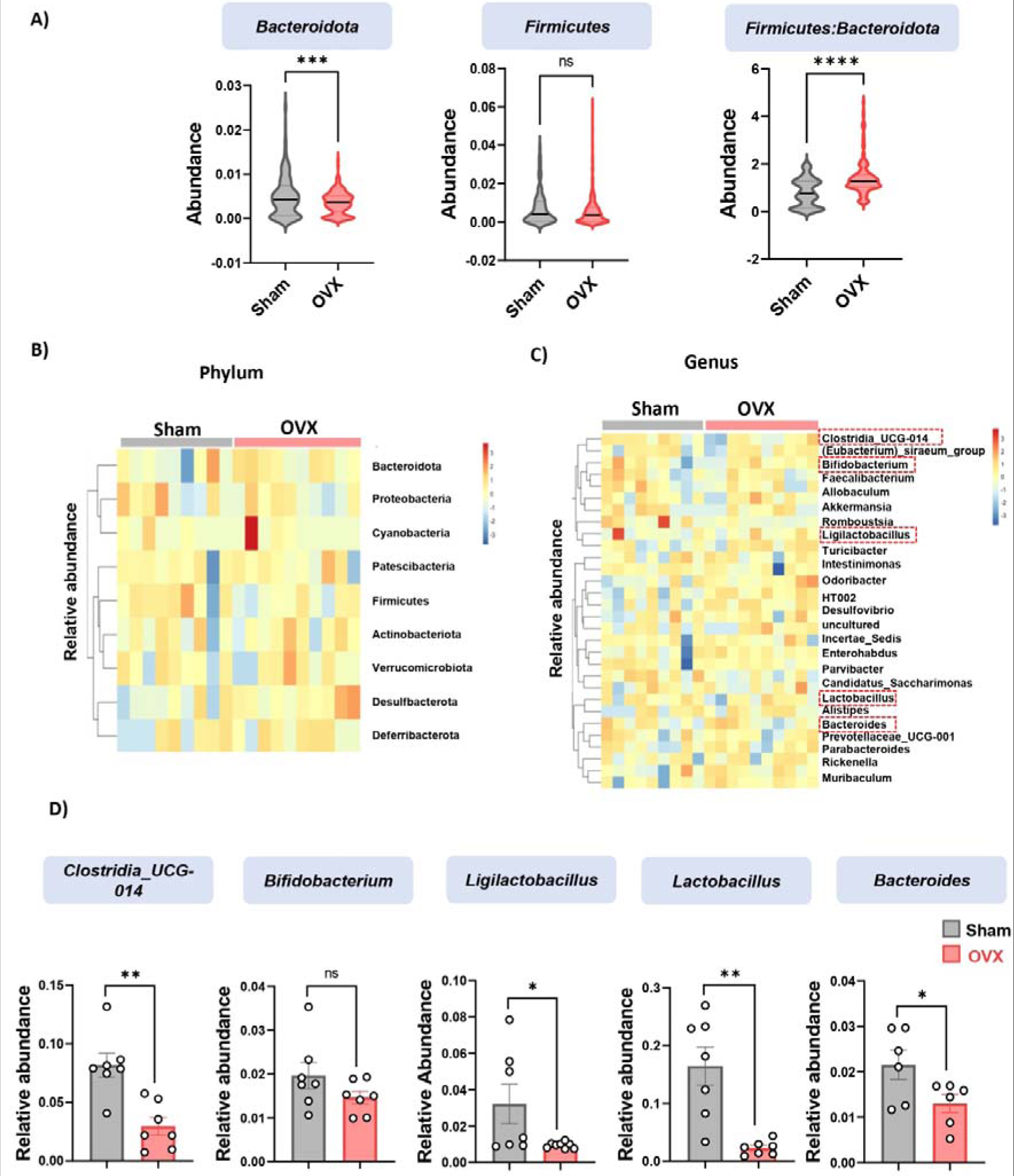
Ovariectomy leads to dysbiosis in mice. A) Plots representing the GM composition at phylum level. B) Heatmap showing differential guts microbiota composition among sham and OVX groups at phylum level. C) Heatmap showing differential gut microbiota composition among sham and OVX groups at genus level. D) Plots representing the GM composition at genus level. Data are represented as mean ± SEM (n = 6-8). Statistical analyses were performed using one-way ANOVA followed by Dunnett’s test. * p < 0.05, ** p < 0.01, and *** p < 0.001.

**Fig. S2:**
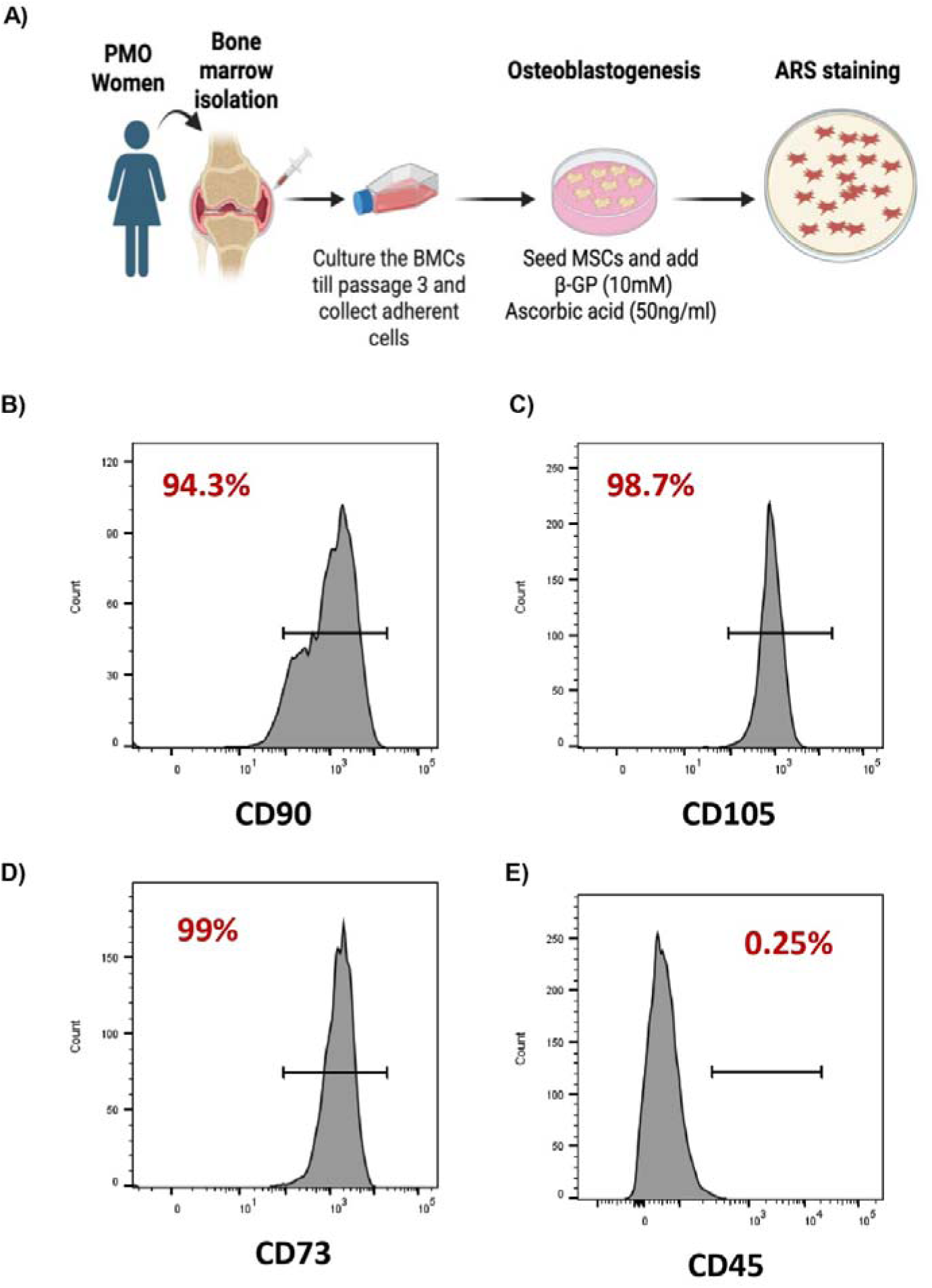
Characterization of Bone Marrow-Mesenchymal Stem Cells. **(A)** Schematic representation of osteoblast culture from human mesenchymal stem cells (MSCs). Flow cytometry analysis of MSCs at passage 3. **(B)** CD90 positive cells, **(C)** CD105 positive cells, **(D)** CD73 positive cells and **(E)** CD45 positive cells.

