## Supplementary material for "Deoxycholic Acid (DCA) a Secondary Bile Acid, Mitigates Ovariectomy-Induced Bone Loss and promotes Bone Healing: A Novel Candidate for Orthopaedic Applications": Table 1

**Table 1:** Primers sequence use in the study.

|  | **Human Primers sequence** | | |
| --- | --- | --- | --- |
| 1 | *CSF1R* | Forward Primer | CTGCTGACTGTTGAGACCTTAG |
|  |  | Reverse Primer | TCCTGCAGAGATGGGTATGA |
| 2 | *DC-STAMP* | Forward Primer | GTGTGCTCCCGCTGAATAA |
|  |  | Reverse Primer | CCCAGATGCAGAGATGGATAAG |
| 3 | *CD47* | Forward Primer | CTGCTCACCAATACCCTGTAAA |
|  |  | Reverse Primer | CGGGAACATCTGATGGACTAAG |
| 4 | *NFATC1* | Forward Primer | TCTGCAACGGGAAGAGAAAG |
|  |  | Reverse Primer | CACGGCTTACGGTTAGAAAGA |
| 5 | *NFkB1* | Forward Primer | CTCCACAAGGCAGCAAATAGA |
|  |  | Reverse Primer | ACTGGTCAGAGACTCGGTAAA |
| 6 | *TRAF6* | Forward Primer | CACGTTGCACACACTCAAATAC |
|  |  | Reverse Primer | CATCTGCGCCTTACCATCTAA |
| 7 | *TREM2* | Forward Primer | CAAGAGACTACTCTGCCTGAAC |
|  |  | Reverse Primer | TCAGAAGTTGTCAGGTGTTCTT |
| 8 | *SYNB* | Forward Primer | AGCCAGCTCTCAAAGGAAATAG |
|  |  | Reverse Primer | GAAGGACTACGGCTGCTAAAG |
| 9 | *CTSK* | Forward Primer | GAAGGGAGTTGGTGTGCTATT |
|  |  | Reverse Primer | GGAGAGAAGCAAAGTAGGAAGG |
| 10 | *ATP6V0D1* | Forward Primer | TGTGCGTGTGTGTGTATGT |
|  |  | Reverse Primer | GTCTCTAAGAGGGTCAGGAGAA |
| 11 | *ATP6V0D2* | Forward Primer | TTACAAGCACCCACCACTAC |
|  |  | Reverse Primer | GAGCTCAGGAGTTCAAGATCAG |
| 12 | *CA2* | Forward Primer | GACTGGCCGTTCTAGGTATTT |
|  |  | Reverse Primer | AGCACTCTTGCCCTTTGT |
| 13 | *SPP1* | Forward Primer | CATATGATGGCCGAGGTGATAG |
|  |  | Reverse Primer | AGGTGATGTCCTCGTCTGTA |
|  | **Mouse Primers sequence** | | |
| 14 | *Tgr5* | Forward Primer | AAGAGCCAAGAGGGACAATC |
|  |  | Reverse Primer | GTAGCTGCTGCTTCCCTAAT |
| 15 | *Fxr* | Forward Primer | GGGCAGAATCTGGATTTGGA |
|  |  | Reverse Primer | GTAGAAGCCCAGGTTGGAATAG |
| 16 | *Nfatc1* | Forward Primer | CCGTCCAAGTCAGTTTCTATGT |
|  |  | Reverse Primer | GTCCGTGGGTTCTGTCTTTAT |
| 17 | *RANKL* | Forward Primer | GGAAGCGTACCTACAGACTATC |
|  |  | Reverse Primer | CTCCCTCCTTTCATCAGGTTAT |
| 18 | *OPG* | Forward Primer | CTTGCCCTGACCACTCTTATAC |
|  |  | Reverse Primer | CTTCCTCACACTCACACACTC |
| 19 | *c-Fos* | Forward Primer | GACCATCTCCGAAATCCTACAC |
|  |  | Reverse Primer | CCCGTCTTGGCATACATCTT |
| 20 | *Osterix* | Forward Primer | CCGAGTCAAGAGTAGGATTGT |
|  |  | Reverse Primer | TAGTGAGCTTCTTCCTGGGT |
| 21 | *Runx2* | Forward Primer | GCTTCTCAGCTTTAGCGTC |
|  |  | Reverse Primer | GTTTCAAGGTGCCGGGAG |
| 22 | *GAPDH* | Forward Primer | AACCCTTAAGAGGGATGCTG |
|  |  | Reverse Primer | AGTTGAGGTCAATGAAGGGG |
| 23 | *Osteocalcin* | Forward Primer | CCAAGCAGGAGGGCAATAA |
|  |  | Reverse Primer | TCGTCACAAGCAGGGTTAAG |
| 24 | *Trap1* | Forward Primer | CACTTTGCACTACAAGACGG |
|  |  | Reverse Primer | TCAGGGGAATGTCCTCACTA |
| 25 | *Ctsk* | Forward Primer | TATGACCACTGCCTTCCAATAC |
|  |  | Reverse Primer | GCCGTGGCGTTATACATACA |
