## Supplementary material for "Deoxycholic Acid (DCA) a Secondary Bile Acid, Mitigates Ovariectomy-Induced Bone Loss and promotes Bone Healing: A Novel Candidate for Orthopaedic Applications": Table 2

**Table 2:** Baseline demographic data of post-menopausal osteoporosis (PMO) subjects recruited for this study.

|  | **Control**  (n= 20) | **PMO**  (n= 57) |
| --- | --- | --- |
| **Age**  **(Years)** | 56.3 ± 9.50 | 63.644 ± 8.54 |
| **Weight (Kg)** | 79.62 ± 11.67 | 55.3 ± 19.49 |
| **T-Score** | 0.14 ± 0.84 | -3.12 ± 0.88**^****^** |
| **BMD** | 0.89 ± 0.42 | 0.70 ± 0.09**^****^** |
